# Argentina Explores Its Bathyal and Abyssal Zone for the First Time Using an ROV: New Biodiversity Discoveries and Unprecedented Public Engagement

**DOI:** 10.64898/2026.05.22.726651

**Authors:** Gregorio Bigatti, Florencia Arrighetti, Graziella Bozzano, Martín I. Brogger, Rodrigo N. Calderón, Nadia Cerino, Ignacio L. Chiesa, Cristina Damborenea, M. Carla de Aranzamendi, Brenda L. Doti, Nahuel E. Farías, Santiago Herrera, Kristen Kusek, Ezequiel Mabragaña, Mariano I. Martinez, Florencia Matusevich, Hannah Nolan, Emiliano H. Ocampo, Leonel I. Pacheco, Guido Pastorino, Pablo E. Penchaszadeh, Sofya Pesternikova, Emanuel Pereira, Jessica Risaro, Renata M. Pertossi, Noelia C. Sánchez, Javier H. Signorelli, Valeria Teso, Diego Urteaga, Johanna N.J. Weston, Jenny Woodman, Daniel Lauretta

## Abstract

Between July 23 and August 12, 2025, members of the scientific group *Grupo de Estudios del Mar Profundo de Argentina* (GEMPA) and collaborators conducted the *Talud Continental IV* expedition in the Mar del Plata Canyon. The expedition was conducted aboard the *R/V Falkor* (*too*) in partnership with Schmidt Ocean Institute (SOI), marking the first deployment of a Remotely Operated Vehicle (ROV) in Argentinean bathyal and abyssal waters. The Mar del Plata Canyon was explored in 2012 and 2013 by CONICET researchers using bottom trawls aboard the *R/V Puerto Deseado* (CONICET, Argentina). This new expedition combined high-definition video surveys, acoustic seafloor mapping, and physicochemical water-column characterization, with *in situ* sensing and sampling of fauna (animal specimens, zooplankton, and environmental DNA), water, sediments, and rock to characterize biodiversity and habitats between 880 and 3900 m. The expedition revealed extensive *Bathelia* cold-water coral reefs, soft-coral gardens, and more than 40 species suspected to be new to science, six of which have already been formally described. Anthropogenic debris, including plastics and fishing gear, was recorded at multiple stations, reaching even the deepest sites, underscoring the extent of human influence on these environments. The *Talud Continental IV* expedition was successful both scientifically and in promoting deep-ocean literacy and engagement, with broad outreach conducted through SOI’s outreach and community engagement programs. The Ship-to-Shore program connected scientists on board with students and educators through live interactive sessions, engaging over 900 students from 19 institutions across Argentina and the United States. The live ROV divestreams, broadcast through SOI’s YouTube and Twitch platforms, reached record levels of public engagement, with ∼19 million total views by July 23rd. The national and international press responded with extensive coverage and interview requests, resulting in over 3,900 international stories. Scientists continued to engage with the public after the expedition through talks at schools and public institutions. The expedition’s achievements promise to usher in a new era of scientific discovery in the Southwestern Atlantic and underscore the value of integrating exploration, conservation, and outreach to inspire wonder and curiosity about the deep-sea in society.

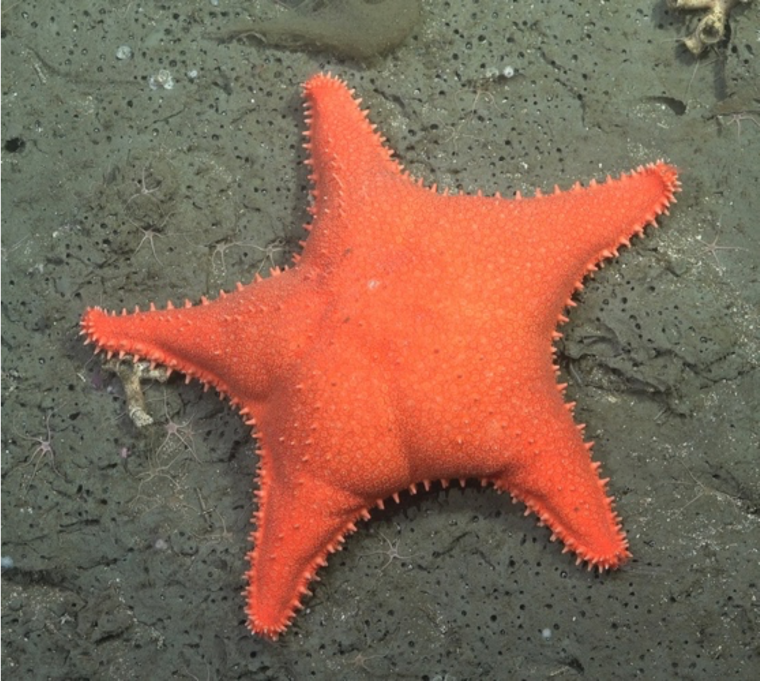

## Introduction

The Southwestern Atlantic continental margin of Argentina remains one of the least-studied oceanic regions in the world, despite its ecological importance and economic potential (Bridges et al., 2023; Bravo et al., 2025). Along this continental margin, many submarine canyons descend to the Argentine basin. Among them, the Mar del Plata Canyon (MdPC) is one of the largest in the South Atlantic Ocean (approx. 38°S; ∼880–3,900 m water depth) (**Fig. 1**). This region lies adjacent to the Brazil-Malvinas confluence (Gordon, 1989; Matano et al., 2010), which may serve as a biogeographic boundary for the deep-sea fauna of the Southwestern Atlantic, marking the northern or southern distribution limit of many taxa.

**Figure 1.**
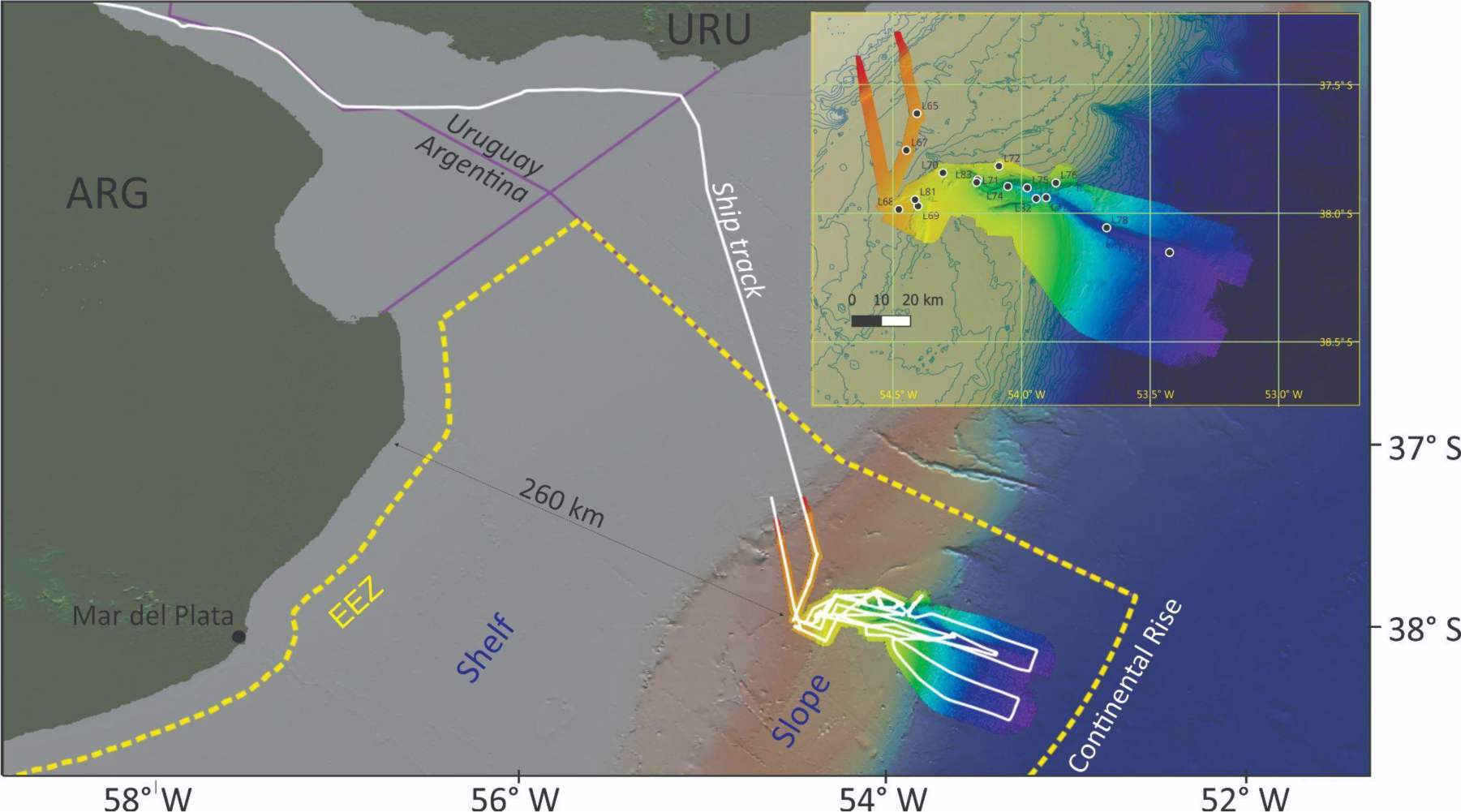
Location of the study area. The international border between Argentina and Uruguay is shown (purple line). EEZ: Exclusive Economic Zone. Base map from GeoMapApp, http://www.geomapapp.org. Inset: Bathymetric map of the Mar del Plata Canyon showing the location (black dots) of ROV *SuBastian* dives. Isobaths every 100 m.

With its 2,000,000 km^2^, the Argentine Continental Margin is one of the largest in the world. It lies between approximately 35°S and 55°S and includes the shelf, the slope, and the continental rise. Several submarine canyons incise the slope, some of them even cutting into the continental shelf. The shelf, covering an area of 960,000 km^2^, extends N-S from the mouth of the Río de la Plata (approx. 37°S) to the southern tip of the Tierra del Fuego Archipelago (approx. 55°S) within water depths that range between 120 and 160 m. Its width ranges from 10 km south of the Isla de los Estados to 850 km at 51°S, opposite the Malvinas Islands (Parker et al., 1996). The shelf edge marks the transition from the relatively flat (slope <1°) continental shelf to the steeper (slope 2°–5°) continental slope. It extends into waters deeper than 200 m until the next change in slope gradient marks the beginning of the continental rise, which is usually found at depths greater than 3500 m (COPLA, 2017).

The MdPC extends from the middle continental slope to the continental rise. It appears to be disconnected from the continental shelf and is therefore classified as a blind, or land-detached, canyon. It probably began to incise into the steep lower continental slope at the end of the Pliocene (Preu et al., 2013). It continued to evolve throughout the Pleistocene and Holocene due to retrogressive erosions and repeated collapses of the walls and headward scarp, which moved further upslope (Krastel et al., 2011; Bozzano et al., 2017).

Previous expeditions in the area relied on dredging and trawling, which provided limited insight into habitat structure and biodiversity. In 2012 and 2013, CONICET researchers planned and conducted three deep-sea biological expeditions to the MdPC area (*Talud Continental I, II*, and *III*), focusing on invertebrate and fish diversity. During those original expeditions, fauna from 64 sampling stations were collected using fishing nets, trawls, and epibenthic sleds, aboard the R/V *Puerto Deseado* (owned by CONICET). Over the past 12 years, more than 400 species have been identified from the material collected during these expeditions, highlighting the region’s remarkable biodiversity.

More than 60 peer-reviewed manuscripts have been published on the fauna of the canyon as a result of those expeditions, and about 30 new species have been described across many groups of invertebrates, including cnidarians (Cerino and Lauretta 2013; Bernal et al., 2018, Risaro et al. 2020), crustaceans (Sganga and Roccatagliata 2016, Doti 2017, Pereira and Doti 2017, Pereira et al. 2019, 2020, 2021, 2023, Roccatagliata 2020, 2023, Chiesa et al. 2024), molluscs (Signorelli and Pastorino 2015, Pastorino 2016a,b, 2019, Pastorino and Sánchez 2016, Teso et al. 2019, 2025, Pacheco et al. 2022, 2024, Sánchez et al. 2023, 2024), echinoderms (Martinez et al. 2014, 2019, Martinez and Penchaszadeh 2017, Rivadeneira et al. 2020, 2025, Flores et al. 2021, 2026, Pertossi et al. 2026), and ascidians (Maggioni et al. 2018). In addition, several species of invertebrates and fishes have been recorded for the first time in the study area: the corals *Fungiacyathus fragilis* (Calderón et al. 2024) and *Dendrobathypathes grandis* (Lauretta and Penchaszadeh 2017); the corallimorpharian *Corallimorphus rigidus* (Lauretta and Martinez 2019); the crinoid *Phrixometra nutrix* (Pertossi et al. 2021), the gastropods *Laubierina peregrinator* (Pastorino 2016a), *Theta lyronuclea* (Sánchez and Pastorino 2020); the valviferan isopods *Acantharcturus brevipleon, Chaetarcturus aculeatus, C. brunneus spinulosus, C. franklini, Fissarcturus granulosus, Litarcturus stebbingi, Dolichiscus georgei, D. marinae* (Pereira et al. 2025); the bony fishes *Bathypterois longipes, Lucigadus nigromaculatus, Paraliparis eltanini, Praematoliparis anarthractae;* and within chondrichthyans, the skate *Bathyraja schroederi* (Vazquez et al. 2016, Cousseau et al. 2020, Mabragaña and Cousseau 2021, Mabragaña et al. 2025). While these efforts yielded valuable biological material, they were insufficient to accurately characterize their spatial distribution and ecosystem characteristics, or to identify human impacts in the area.

Building on the known biological importance of this canyon, the GEMPA (*Grupo de Estudios del Mar Profundo de Argentina)* focused on improving the ecological understanding of the canyon through a fourth expedition. The *Talud Continental IV* expedition, “Underwater Oases of the Mar del Plata Canyon”, FKt250712, conducted from July 23 to August 12, 2025 aboard the R/V *Falkor (too)* in collaboration with the Schmidt Ocean Institute (SOI), aimed to (1) provide the first *in situ* visual characterization of deep-sea habitats in the Mar del Plata Canyon, (2) collect specimens and environmental samples to document biodiversity, (3) characterize the seafloor that host the different habitats with a focus on vulnerable marine ecosystem indicator taxa, (4) assess the reach of anthropogenic impacts, and (5) engage the public through ship-to-shore activities with schools and real-time talks of deep-sea exploration.

## Materials and Methods

The *Talud Continental IV expedition* combined high-definition video surveys, acoustic seafloor mapping, physicochemical water-column characterization, in situ sensing and sampling of fauna (animal specimens, zooplankton, and environmental DNA), water, sediments, and rock geology, to characterize deep-sea biodiversity and habitats between 880 and 3900 m.

The scientific team aboard the ship consisted of 27 researchers: 26 biologists, mostly with taxonomic specialties, and one geologist. Of the 26 biologists, 24 were based at Argentine institutions, while the remaining two work at foreign institutions (see co-author affiliations). The vessel crew of the R/V *Falkor (too)* included officers (captains, chief officers, and second and third officers), engineering staff (chief engineers and second and third engineers), technical staff (marine technicians, AV/IT engineers, and electro-technical officers), deck and bosun personnel (bosuns, fitters, and able bodied seamen), support services (pursers, galley stewards, and chefs), and specialized ROV and multimedia engineers and technicians. This crew ensured safe operations, efficient coordination of ship and ROV activities, and high-quality technical support for scientific and outreach objectives. All outreach objectives were further supported by SOI’s shoreside communications team.

### Ship-Based Exploration

Seafloor morphology was characterized using a suite of high-resolution Kongsberg acoustic systems, including the EM124 for deep-water mapping, the EM712 for intermediate depths, and the EM2040 MK II for shallow-water environments. Sub-surface sedimentary structures were resolved using a Kongsberg SBP29 sub-bottom profiler. Water-column currents were monitored using Teledyne RDI OS38 (38 kHz) and WH300 (300 kHz) Acoustic Doppler Current Profilers (ADCPs).

The vessel’s underway seawater system continuously monitored sea surface parameters using a WET Labs C-Star transmissometer, a WET Labs ECO FL fluorometer, and a Sunburst Sensors AFT-pH sensor. Thermosalinograph data were collected via Sea-Bird Electronics model 45 and SBE-38 sensors. Meteorological data were recorded by an Eppley SPP/PIR radiometer, Gill Windsonic anemometers, Paroscientific Met4 stations, and Gill Instruments MaxiMet Marine GMX560 weather stations. On-board laboratory facilities included Leica EZ4 W and Leica S Apo stereo microscopes, a 4 °C cold room, and ultra-freezers (-30 °C and -80 °C) for specimen preservation.

### ROV-Based Exploration

Deep-sea exploration was executed using the ROV *SuBastian*, a custom-built work-class vehicle rated to 4,500 m. Scientific navigation was provided by Sonardyne WMT (forward) and WSM6 (aft) Ultra-Short Baseline (USBL) beacons. Environmental data at the bentho-pelagic interface were captured using a Sea-Bird SBE 49 FastCAT CTD and an Aanderaa 4831 oxygen optode. Precise depth and pressure measurements were obtained via Valeport Mini IPS and Paroscientific Digiquartz 8CB7000-1 sensors. Visual data were recorded using a SULIS Subsea Z70 4K science camera with 10X zoom and a Z71 HD situational camera. Some of these instruments are shown in **Fig. 2**. Species annotations were logged in real-time using SeaLog software v2.x. During the expedition, the ROV *SuBastian* was operated by a team of eight SOI technicians, working in pairs on two-hour shifts under the guidance of the scientific team.

**Figure 2.**
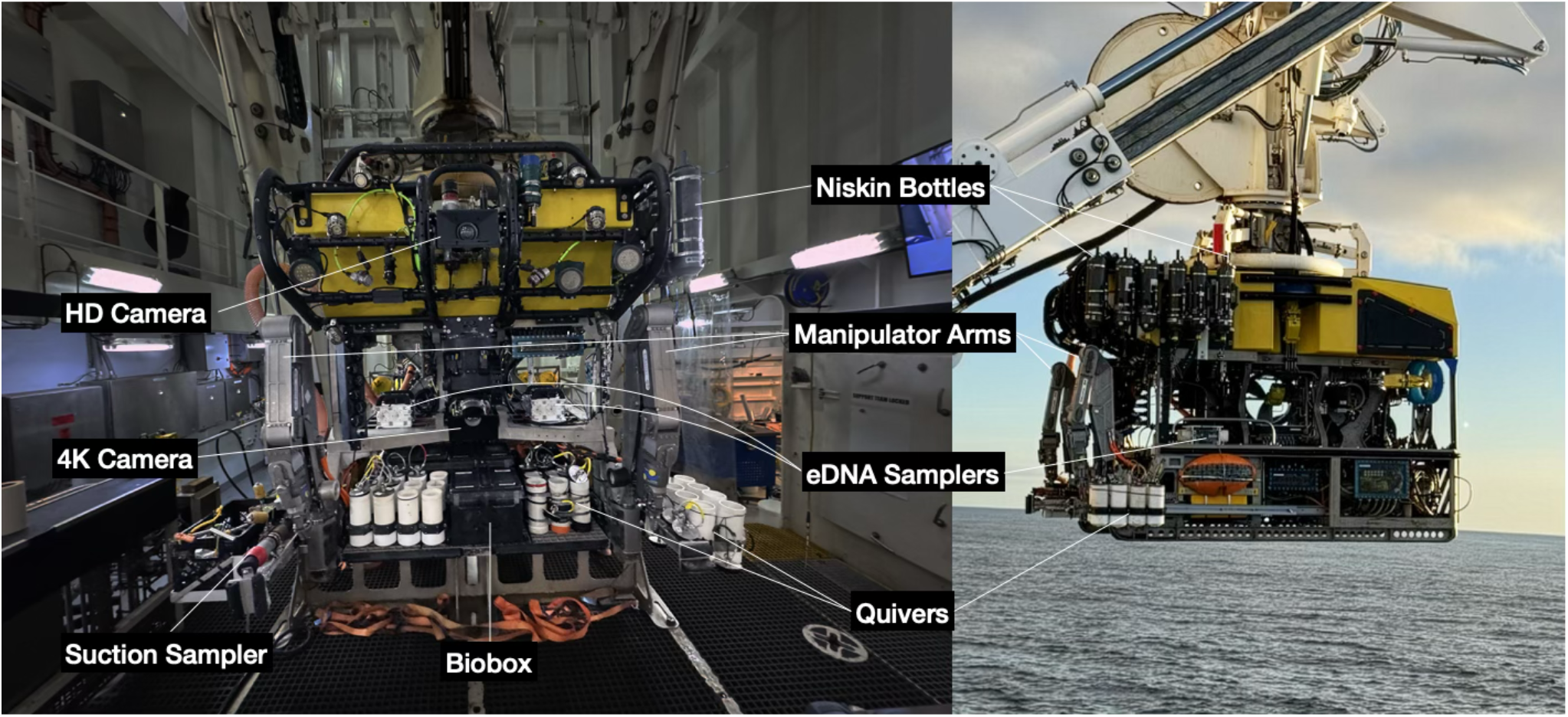
ROV *SuBastian* with tools and instruments used on expedition *Talud Continental IV* onboard the R/V *Falkor (too)*. Photo credits: Santiago Herrera.

Each dive included standardized 100 m transects, high-definition video recording, species annotations using the SeaLog software, bottom mapping, and targeted sampling of fauna, sediments, and water. The imagery data were field-annotated by all scientists on board, based on their taxonomic expertise. On land, all specimens are being identified to the lowest possible level. Each raw video will be edited to remove unusable sequences and used to estimate the abundance of different species. Substrate type was recorded, along with the percent cover of colonial or encrusting fauna, following the methods of Gasbarro et al. (2022). The analysis of these data is ongoing, and results will be reported in subsequent publications describing the distribution and abundance of vulnerable marine ecosystem indicator taxa at each site.

### Biological Sampling: Animal Samples

Benthic fauna samples were collected using *SuBastian*’s two seven-function T4 Schilling hydraulic manipulator arms to place them into bioboxes and quivers with lids (Fig. 3). Faunal samples were also collected using a suction sampler equipped with seven 2.3 L sampling containers with various mesh sizes.

**Figure 3.**
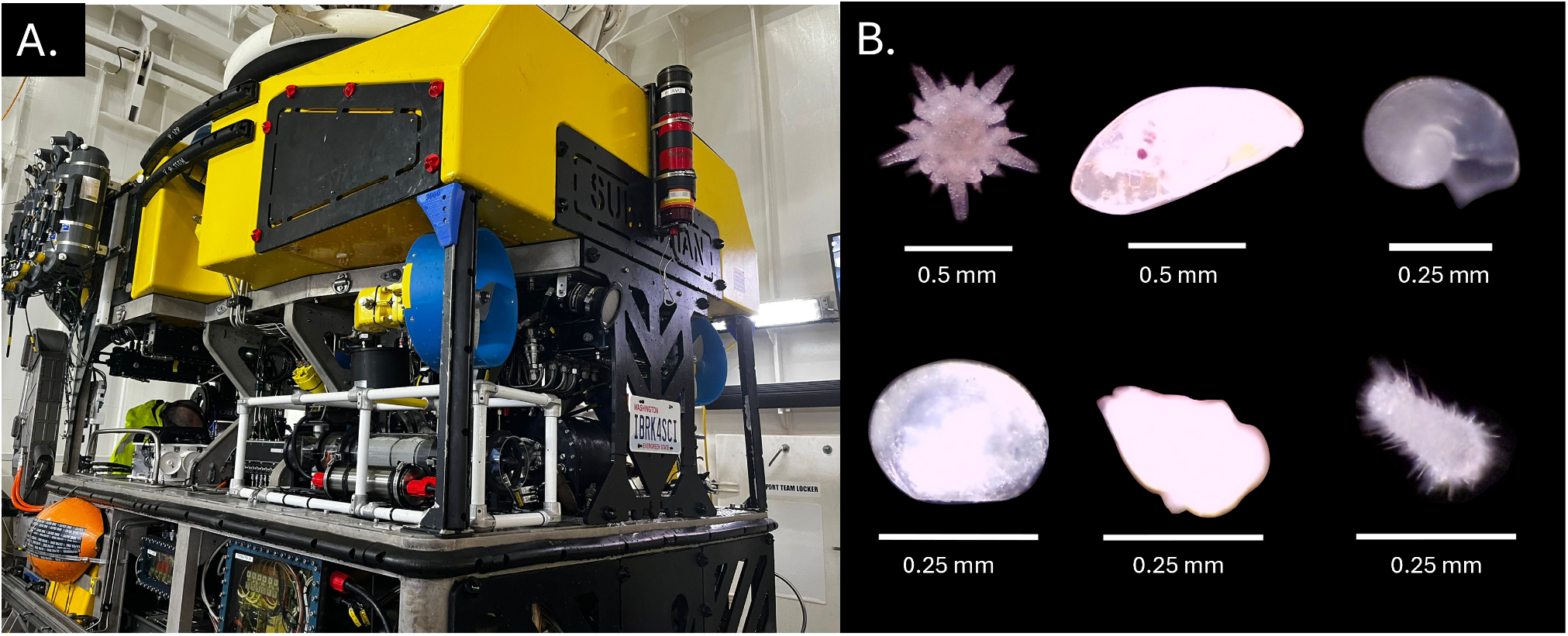
Zooplankton sampling. (A.) *DeepZoo* mounted to ROV *SuBastian*. (B.) Six representative morphotypes collected by *DeepZoo*. Top left to bottom right: ophiuroid, barnacle cyprid, gastropod, bivalve, cnidarian planula, and aplacophora. Photo credits: Johanna Weston.

Once on board, the specimens were immediately labeled with a unique collection number. Tissue subsamples were preserved for molecular work in 96–100% ethanol, RNA later, and/or frozen at -80°C. Most specimens were fixed and preserved in 96% ethanol, but some groups required fixation in 4% seawater-buffered formalin (e.g., sea anemones, ascidians, fishes, and large holothurians). For specimens considered potentially new, identifications will follow standard morphological and DNA sequencing procedures, including stereoscopic microscopy, scanning electron microscopy, morphometry, and histology. The collected data will be made available as soon as possible by publishing manuscripts and submitting datasets to open online repositories (e.g., GenBank for genetic data, OBIS for biodiversity data). The collected specimens will be identified, curated, and deposited in the national invertebrate collection at the MACN to ensure long-term preservation and accessibility to the scientific community.

### Biological Sampling: Environmental DNA

Environmental DNA (eDNA)-containing particles were captured using two Multipuffer (Aquatic Labs) high-volume *in situ* filtration systems mounted on the front of *SuBastian*, featuring eight independently addressable pumps and integrated temperature and pressure sensors. High-volume filtration is critical for representative sampling in the deep-sea (Herrera et al. 2026). Filtered water samples were used to capture traces of DNA shed by marine organisms. Metabarcoding the resulting eDNA enables rapid biodiversity surveys and accurate taxonomic identification of the communities present at the time *SuBastian* visited each site, including cryptic or rare taxa that visual surveys miss. The power of eDNA observations relies on the taxonomically curated reference sequences that will be produced from the collected animal specimens. For this expedition, the *in situ* filtration systems were outfitted with 90-mm-diameter PES filters housed in custom-designed, reusable cartridges for high-flow, medium-to high-volume eDNA sampling in the deep sea.

### Biological Sampling: Zooplankton

During the expedition, zooplankton, both meroplankton and holoplankton, were collected using *DeepZoo*, a novel autonomous sampler. Understanding the diversity and dispersal of benthic invertebrate early-life stages is key to studying population distribution and connectivity across deep-sea habitats. Developed at WHOI’s AVAST Innovation Hub, *DeepZoo* is a cost-effective, lightweight (∼7 kg in water) all-depth device currently at the working-prototype stage (**Fig. 3A**; Weston et al., in review). It integrates two systems: (1) sample collection, composed of a gasketed lid with an oil-filled 10 RPM gear motor (ServoCity, USA), a 63 µm Nitex mesh net, and a T200 thruster (Blue Robotics, USA) to drive water flow; and (2) system control, housed in titanium and containing electronics, including a 14.8 V, 10 Ah rechargeable lithium-polymer battery (Blue Robotics). Pre-programmed in Arduino IDE 2.3.6 (Arduino 2024), *DeepZoo* runs autonomously, controlling motor and thruster operation according to dive plans. The two systems are contained within a lightweight PVC frame with SpeedRail connectors and a wire-mesh base (91 × 28 × 30 cm). Upon recovery, the net tube was submerged in chilled, filtered seawater (0.5 µm) and processed in a 4 °C cold room. Samples were rinsed and sieved through a 63 µm mesh. Sorting targeted both overall diversity (one individual per morphotype) and benthic diversity (all larvae and early-life stages). Picked organisms were transferred to 6-well plates, imaged on either a Leica S Apo (Flexcam C3) or an Amscope DM300HD compound microscope (brightfield/darkfield, 4x–10x) depending on size, and individually preserved in 0.5 mL or 1.5 mL tubes in ethanol (96%) for downstream analyses such as DNA barcoding. Whole residual samples were also preserved in ethanol. All samples are archived at the Invertebrate Collection of the MACN.

### Biological Sampling: Lander

Biological sampling was also conducted using the SOI Lander (**Fig. 4**), which was equipped with multiple baited traps of varying sizes designed to collect crustaceans and fish. The baited traps included: (1) three PVC tube traps with a funnel on one end and 300 micron mesh on the other side, aimed at some small crustaceans like amphipods; (2) one canister with side punctured hole also aimed at small crustaceans; (3) one large rectangular mesh cage (36 cm wide, 36 cm high, 71 cm long) with one entrance for crustaceans or fishes; (4) one cylinder mesh cage for crustaceans or fishes; and (5) one bespoke large net trap with an entrance at the top and a net that draped down to the seafloor and targeted crabs. The traps were baited with several types of bait, including canned mackerel, canned liverwurst, and whole fish. When on deck, baited-trap specimens were placed in chilled seawater and immediately identified by a unique collection number. Specimens were subsampled for molecular analysis and frozen at -80°C. Most of the specimens were fixed and preserved in 96% ethanol. The lander was deployed three times with baited traps and outfitted with *DeepZoo*, with a descent rate of ∼50 m/min and an ascent rate of ∼15 m/min. Lander deployments 1 and 3 were visited on the seafloor with *SuBastian* to obtain visual confirmation of adequate positioning and instrument functionality.

**Figure 4.**
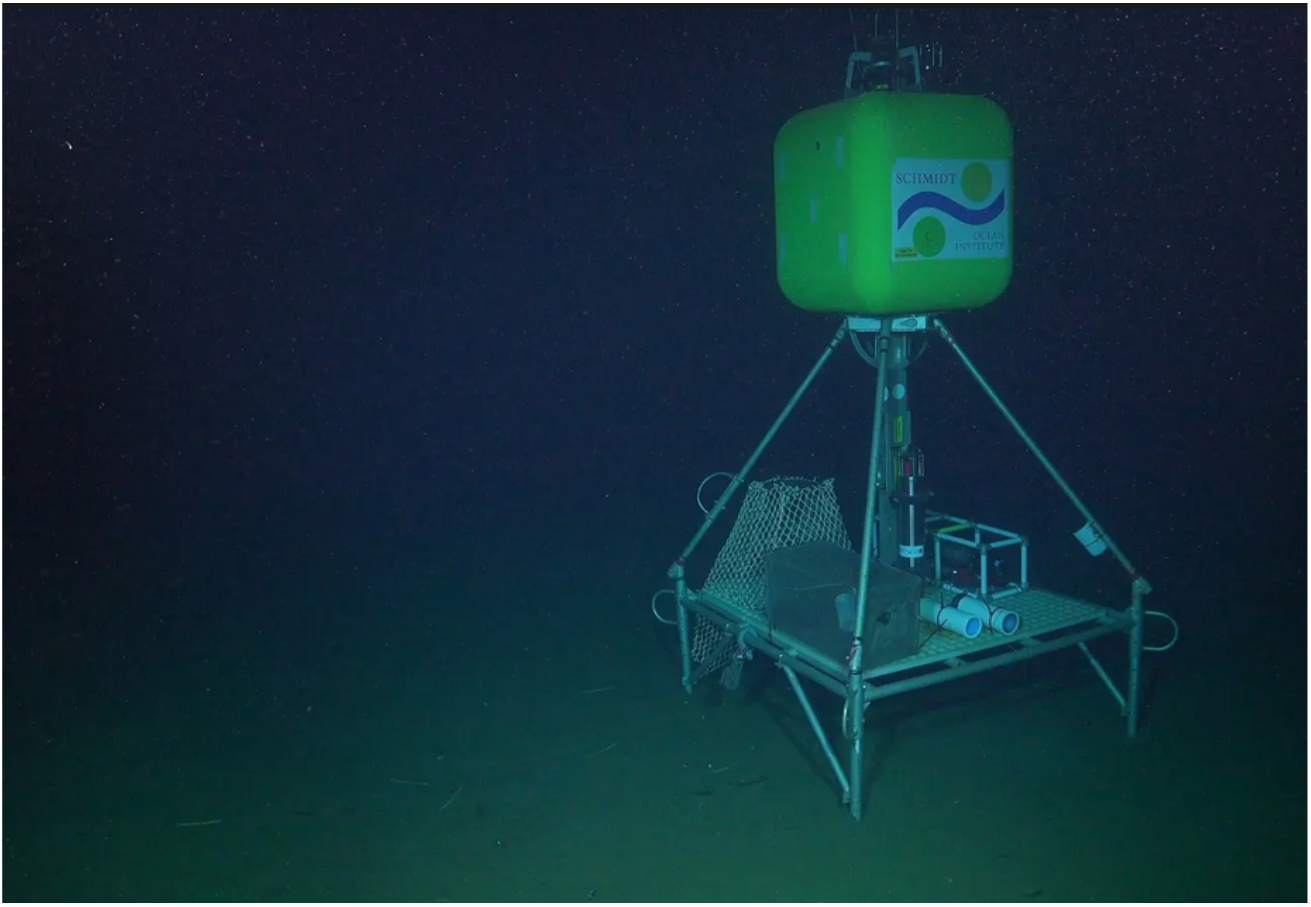
SOI Lander outfitted with multiple sizes of baited traps and *DeepZoo* during deployment 1 to 878 m. Photo credit: Schmidt Ocean Institute.

### Sediment Sampling: Grain Size and Carbon Cycling

Sediment samples were collected using the 27-cm-long push cores or sediment bags installed on the ROV *SuBastian*. Rocks that served as substrate for organisms were also obtained when possible. Sediment slices from additional samples are being analyzed to estimate carbon stocks (both organic and inorganic) and burial rates. Organic matter will be estimated from burned (450 °C) and unburned sediment. Analyses of stable carbon and nitrogen isotopes (δ^13^C and δ^15^N) will help identify the main sources of organic matter. At the same time, ^210^Pb-based geochronology, with supported ^210^Pb estimated via ^214^Pb, will be used to estimate sediment accumulation rates and thereby organic-carbon burial rates.

### Anthropogenic Impact Assessment: Microplastics & : Marine Litter

Seawater, sediment, and animal tissues were sampled to assess the existence of microplastics in the MdPC. Water samples were obtained via 5-liter Niskin bottles mounted on *SuBastian’s* frame and triggered from the ROV control room. After recovery, water samples were transferred into clean bottles and frozen at -20°C until analysis. Sediment samples were collected using push-cores. On board, only the upper 0–5 cm layer of sediment was used for analysis. The samples were frozen at -20°C until analysis. Tissue samples from the most abundant species were identified and placed individually in plastic bags, then frozen at -20°C for preservation before analysis. Microplastic analyses are ongoing and include microscopic examination of microplastics in each sample to quantify their abundance, and Raman spectroscopy to determine their chemical composition.

Anthropogenic debris, or marine litter, was recorded on video and, when possible, collected using the *SuBastian*. For each dive, video recordings are being independently reviewed to document the geographic coordinates (latitude and longitude) and depth of each piece of anthropogenic debris.

### Outreach: Artist-at-Sea

On each expedition, SOI includes artists as part of its Artist-at-Sea program. The program is designed to foster meaningful dialogue between scientists and artists, who then channel the inspiration from the at-sea experiences to produce art that inspires new understanding about, and connection to, the ocean. During this expedition, Pablo E. Penchaszadeh, the Artist-at-Sea, was inspired by the seafloor images to paint textures and transparencies using diverse substrates, media, and techniques (**Fig. 5**). Pablo is also a marine biologist, a senior scientist at CONICET, and a mentor to many in the expedition’s scientific group.

**Figure 5.**
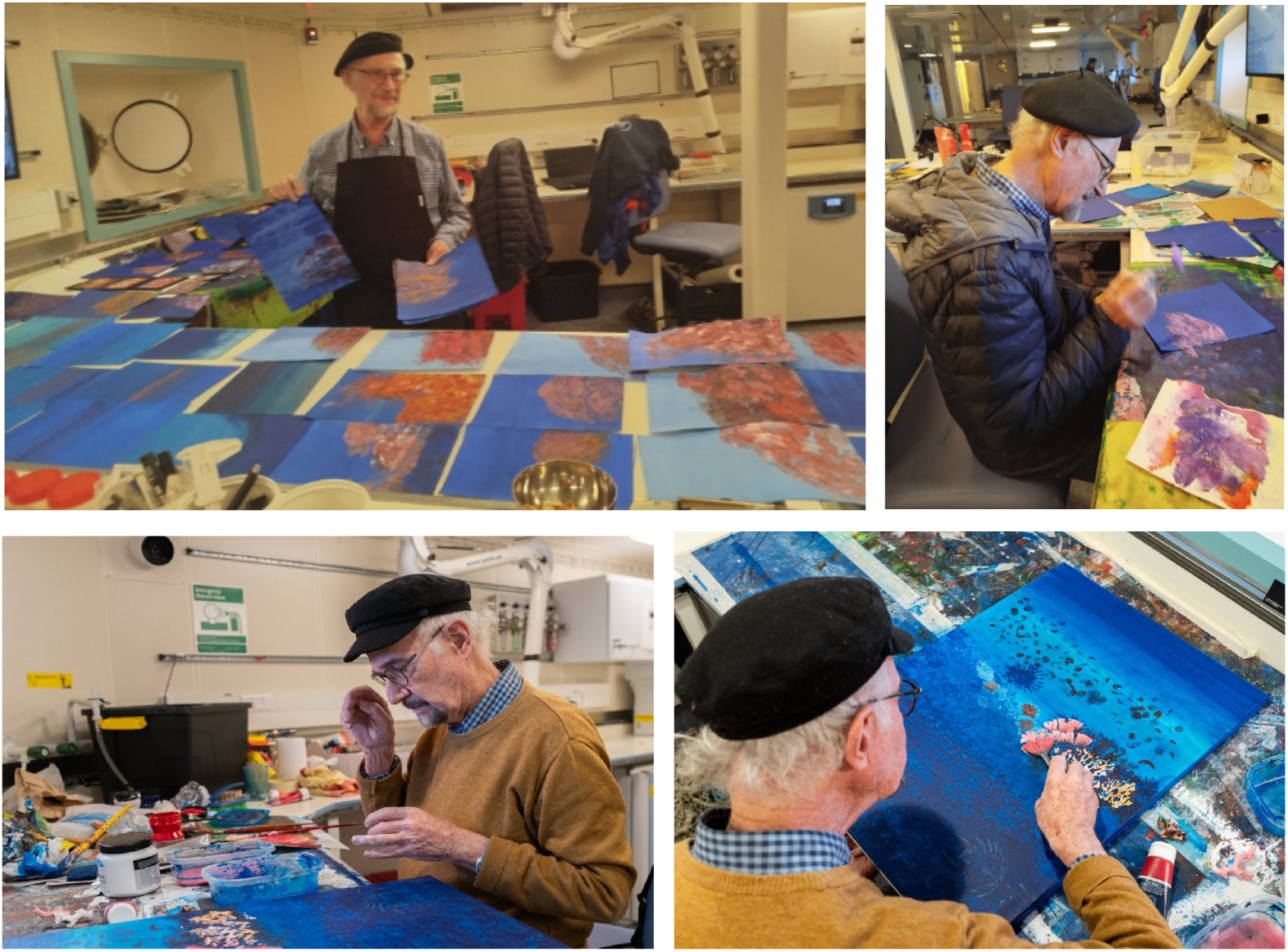
The Artist-at-Sea, Pablo Penchaszadeh, painting during the *Talud IV* expedition. Photo credits: Gregorio Bigatti and Misha Vallejo Prut/Schmidt Ocean Institute.

### Outreach: Ship-to-Shore Program

The Ship-to-Shore program is a regular educational SOI initiative that connects scientists aboard the R/V *Falkor (too)* with the general public, and prioritizes students and educators from the country where the expedition takes place. We conducted 19 ship-to-shore sessions during the expedition, six of which received funding through SOI’s technology support small grants program.

Each session was conducted online via Zoom and guided by one of the onboard scientists, who led participants through key areas of the vessel, including the bridge, laboratories, and the ROV control room, thereby creating an interactive and conversational experience. The program is supported by the onboard multimedia technician, who manages the equipment, camera, schedule, and connection. Other scientists and crew members joined the sessions spontaneously to explain their work and answer questions. The sessions typically lasted 45–60 minutes and offered students a rare opportunity to interact directly with a working oceanographic research team in real time. Although a general plan is in place for all ship-to-shore activities, interactions with the public and researchers made each activity unique.

### Outreach: Divestreaming of ROV Operations

All ROV dives were broadcast live through the Schmidt Ocean Institute’s YouTube and Twitch streaming channels, and promoted through the institute’s social media channels. During each dive, members of the science team described the ongoing exploration, identified species and geological features, and answered public questions submitted via the streaming live chats.

### Outcomes

The 21-day expedition (July 23 – August 12, 2025) achieved a total ROV *SuBastian* dive time of 291 hours, 24 minutes, and 31 seconds. Of these, 223 hours were spent on the bottom. Of the 18 ROV deployments, 17 were successful scientific dives (Dive SO824 was aborted during descent to the seafloor at 359 m due to a technical issue), resulting in 1035 sampling events. The maximum depth reached during the expedition was 3997 m during dive SO821 (**Table 1**). The duration of the dives and on-bottom broadcasts varied considerably, ranging from 4-5 hours to more than 24 hours. To sustain continuous operations, 24 hours/day, scientists were divided into two 12-hour shifts.

**Table 1.** ROV *SuBastian* dive stations during the *Talud Continental IV* expedition (FKt250712) to the Mar del Plata Canyon within the Argentine EEZ. Dive SO824 was aborted and is not included. Latitude, longitude, and depth represent average values (dive centroid) calculated from the on-bottom data for each dive.

| ROV Dive | Off Deck Time (UTC) | Bottom Duration (h) | Lat. (dd) | Lon. (dd) | Avg. Depth (m) | Track Length (km) | Sampling events |
| --- | --- | --- | --- | --- | --- | --- | --- |
| SO809 | 25-07-24 23:11:19 | 03:44 | -37.61131 | -54.40544 | 900 | 0.66 | 58 |
| SO810 | 25-07-25 09:25:32 | 05:01 | -37.75753 | -54.44767 | 876 | 1.34 | 48 |
| SO811 | 25-07-26 01:45:06 | 05:17 | -37.98530 | -54.47607 | 1013 | 1.07 | 32 |
| SO812 | 25-07-26 12:57:03 | 07:45 | -37.97174 | -54.40752 | 1203 | 4.92 | 31 |
| SO813 | 25-07-27 11:40:07 | 04:16 | -37.84353 | -54.30611 | 1262 | 2.24 | 17 |
| SO814 | 25-07-27 23:22:01 | 12:34 | -37.86939 | -54.16692 | 1915 | 3.14 | 62 |
| SO815 | 25-07-28 19:58:22 | 07:34 | -37.81815 | -54.09282 | 1417 | 2.17 | 27 |
| SO816 | 25-07-29 11:34:21 | 18:01 | -37.88604 | -54.05982 | 2243 | 5.39 | 58 |
| SO817 | 25-07-30 13:16:24 | 16:35 | -37.89399 | -53.97079 | 2328 | 4.57 | 54 |
| SO818 | 25-07-31 13:25:42 | 19:29 | -37.89394 | -53.87838 | 2380 | 5.12 | 56 |
| SO819 | 25-08-01 16:19:33 | 24:19 | -37.94274 | -53.90342 | 3041 | 5.92 | 114 |
| SO820 | 25-08-04 09:30:43 | 20:48 | -38.05144 | -53.66670 | 3582 | 3.71 | 94 |
| SO821 | 25-08-05 15:59:44 | 09:51 | -38.15139 | -53.42647 | 3780 | 8.34 | 50 |
| SO822 | 25-08-06 20:04:02 | 21:04 | -37.94816 | -54.41308 | 1079 | 5.39 | 92 |
| SO823 | 25-08-07 23:02:35 | 13:26 | -37.94245 | -53.93678 | 2893 | 3.93 | 62 |
| SO825 | 25-08-09 02:00:21 | 18:50 | -37.87877 | -54.17670 | 1888 | 3.98 | 94 |
| SO826 | 25-08-10 03:42:46 | 14:15 | -38.01843 | -54.50397 | 984 | 1.74 | 86 |
| TOTALS |  | 222:49 |  |  |  | 63.64 | 1035 |

### Biodiversity and Habitat Characterization

The biological communities encountered were heterogeneous and correlated with the canyon’s different environments (walls vs. valley, deep vs. shallow areas). Extensive cold-water coral reefs, along with large fields and gardens of soft corals, and abundant echinoderms, fish, nemerteans, crustaceans, mollusks, and other organisms, were observed and collected (**Fig. 6**). Benthic assemblages displayed high diversity and structural complexity, suggesting the canyon acts as a biodiversity hotspot in the Argentinian margin.

**Figure 6.**
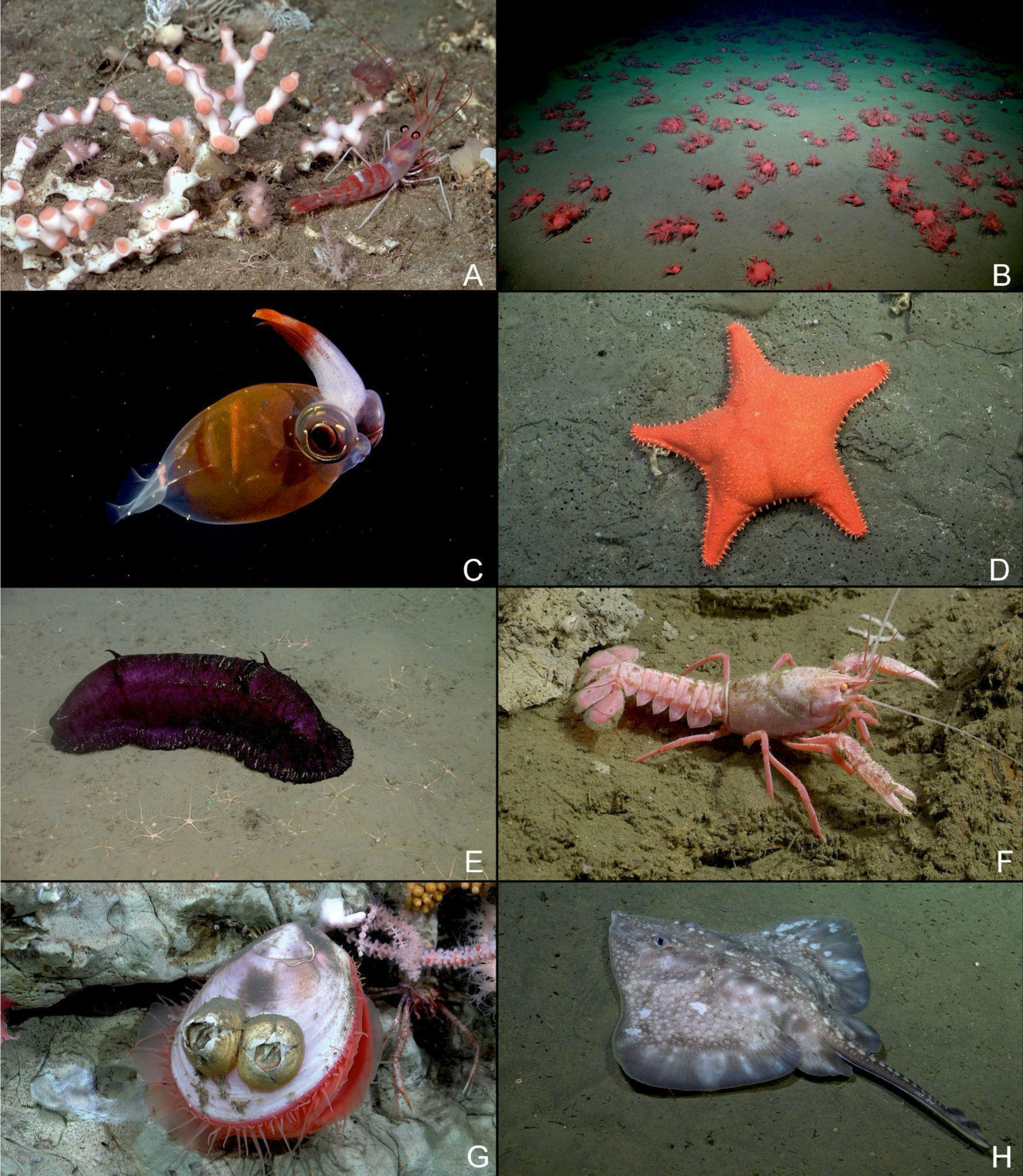
Examples of species found during the expedition. A. Reef-forming cold water coral *Bathelia candida* Moseley, 1880 (Hexacorallia: Oculinidae). B. Fields of *Heteropolypus* sp. (Octocorallia: Coralliidae). C. Crystal squid of the family Cranchiidae (Mollusca: Cephalopoda). D. Sea star *Hippasteria phrygiana* (Parelius, 1768) (Asteroidea: Goniasteridae). E. Elasipodid sea cucumber *Benthodytes violeta* Martinez et al. 2014 (Holothuroidea: Psychropotidae). F. Patagonian lobsterette *Thymops birsteini* (Zarenkov & Semenov, 1972) (Malacostraca: Nephropidae). G. Field clams *Acesta* sp. (Bivalvia: Limidae) associated with barnacles *Tetrachaelasma southwardi* Newman & Ross, 1971 (Cirripedia, Balanomorpha, Bathylasmatidae). H. The thickbody skate *Amblyraja frerichsi* (Krefft, 1968) (Elasmobranchii: Rajidae). Images courtesy of the Schmidt Ocean Institute.

Evident differences in community structure were observed over the steep North and South face in contrast to the plain canyon axis. In addition, in some stations, key/dominant species such as *Heteropolypus “remolachas”*, or *Bathelia candida* Moseley, 1880, were documented. Over 400 species of invertebrates were identified, where corals and echinoderms were usually dominant. For fish, more than 50 species were recognized, although in some cases they were not identified to species because few specimens were collected. The most commonly observed families were Macrouridae Bonaparte, 1831, and Moridae Moreau, 1881, and the Anguilliformes order. Within the skates, almost all the known deep-sea species of the area were observed.

Overall, over 40 species were identified as potentially new to science. These include corals, anemones, sponges, platyhelminths, nemerteans, mollusks, crustaceans, echinoderms, ascidians, and fishes. Descriptions based on morphological and genetic data are being conducted to confirm the taxonomic status of the collected material. Sediment analysis is ongoing, and the results will be reported in subsequent publications. So far, six new species, one crinoid and five bryozoans, have been formally described (Pertossi et al. 2026, López Gappa & Urteaga 2026).

### Environmental DNA

Across all dives, the system collected 104 discrete eDNA samples, totaling over 26,000 liters. The pumps were controlled in real-time from the surface, with operators using the ROV camera feeds to determine when to activate or pause sampling. This setup ensured the collection was targeted to scientific objectives and avoided contamination during sediment resuspension events. This technology represents an innovative advance in deep-sea biodiversity monitoring, allowing habitat-aware, volume-adequate sampling (Herrera et al. 2026). Laboratory analysis of collected material is ongoing. Results will be reported in subsequent publications. The successful first deployment of these systems on *SuBastian* demonstrated the utility of this approach for real-time, targeted environmental sampling, setting the stage for expanding their use in future campaigns.

### Zooplankton

*DeepZoo* was deployed nine times in a horizontal configuration—three on the SOI Lander and six on *SuBastian* (**Fig. 3A and 4**). It was programmed to sample only while the vehicle was estimated to be on the bottom. Sampling durations averaged 692 minutes (range: 320–960 minutes). The cumulative sampling duration was 104 hours, spanning habitats from the continental shelf above the canyon (878 m; Lander 1/SO810) to beyond the canyon mouth (3739 m; Lander 3/SO821). In total, 142 morphotype tubes were prepared and 298 specimen images taken. Samples represented at least 19 taxa across Annelida, Cnidaria, Mollusca, Echinodermata, Arthropoda, Chaetognatha, Foraminifera, and undetermined Animalia (**Fig. 3B**). Zooplankton analysis is ongoing, and the results will be reported in subsequent publications.

### Sediment

During the 17 successful *SuBastian* dives, 27 push cores were retrieved for sedimentological purposes, with 1 or 2 cores per dive. Each core was immediately subsampled at specific intervals (0–1 cm, 1–2 cm, 4–5 cm, 9–10 cm, 14–15 cm, 19–20 cm, bottom), and the sediment was stored at 4 °C. Grain-size analyses will be performed using a CILAS 1190 laser particle size analyzer, covering a range of 0.04-2,500 μm, thereby capturing grain-size distributions from clay to sand. The objective was threefold: to characterize the seabed lithology, to interpret the sedimentary processes operating within the canyon domain, and to determine the substrate type best suited to each deep-sea habitat.

Overall, the sediment samples will provide a solid baseline to help identify ecologically important areas with high carbon storage potential and to improve our understanding of the role deep-sea sediments play in regional carbon dynamics and the broader carbon cycle. Sediment analysis is ongoing, and the results will be reported in subsequent publications.

### Anthropogenic Debris

Debris such as plastics, fishing gear, and footwear was observed at multiple depths, including sites below 3,000 m. These observations confirm that even remote deep-sea habitats are not insulated from human influence. The origin of the debris is not clear at the moment. Video analysis is ongoing, and the results will be reported in subsequent publications.

### Public Engagement and Outreach

The expedition’s outreach achieved unprecedented success. Livestreamed dives attracted record-breaking public engagement. From July 23 to August 12, 2025, Schmidt Ocean Institute’s YouTube channel (https://www.youtube.com/@SchmidtOcean) accumulated 19 million views and ∼460,000 new subscribers (net change from a pre-expedition baseline of ∼59,000 subscribers) (**Fig. 7**). The average daily views were ∼814,000. Approximately 82.5% of the audience was located in Argentina. Peak concurrent viewership exceeded 92,000. Engagement included schools, community centers, museums, and media platforms. This expedition was the institute’s most widely viewed ocean exploration livestream to date, and the largest ocean science outreach initiative in Argentine history. In the words of Wendy Schmidt, “Wonder is part of human nature, and it’s easy to spark, when we know where to look” (Schmidt, 2025).

**Figure 7.**
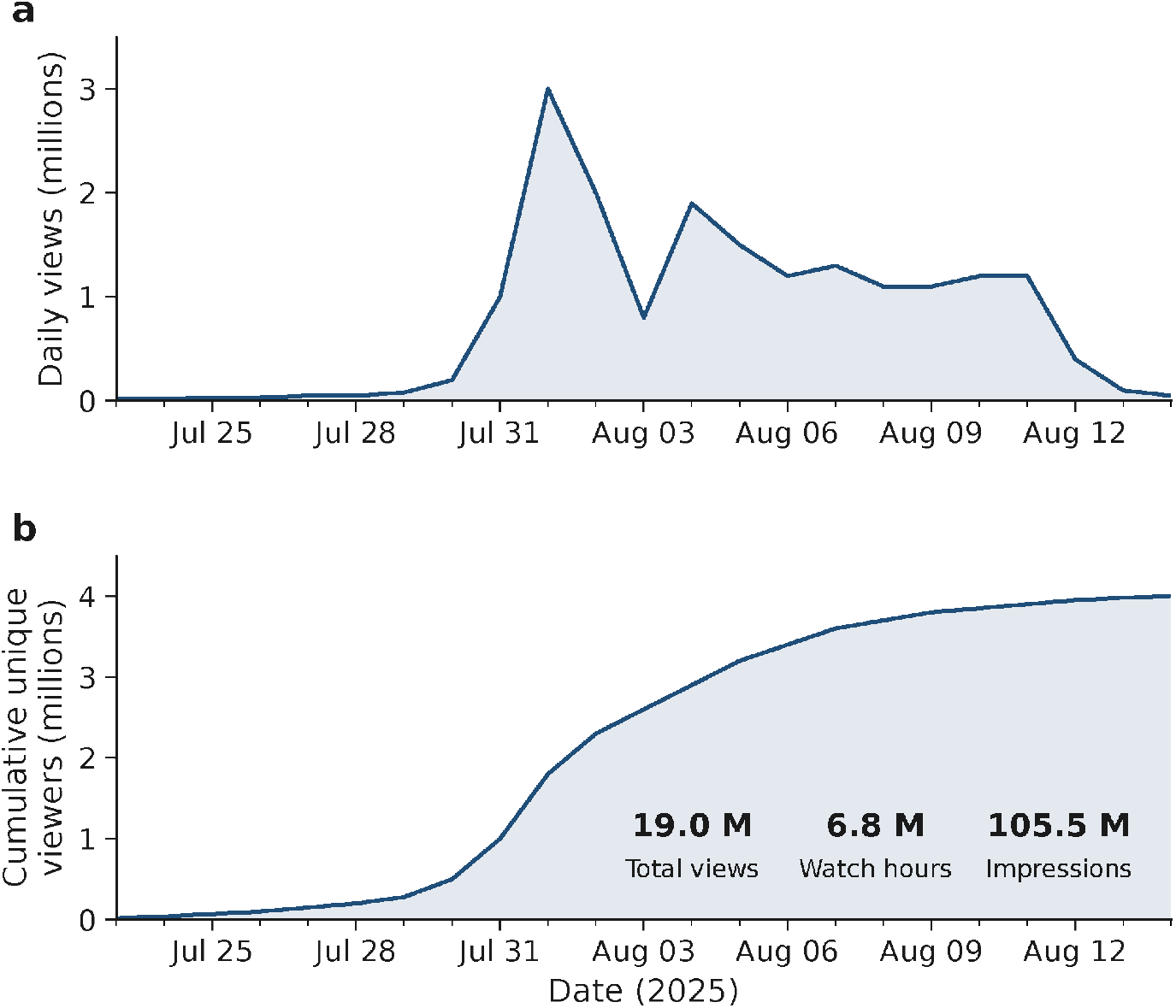
Schmidt Ocean Institute YouTube channel engagement metrics during the 23-day window of 23 July – 14 August 2025, encompassing the *Talud Continental IV* expedition aboard R/V *Falkor (too)*. (**a**) Daily views (millions), peaking at 3.0 million on 1 August. (**b**) Cumulative unique viewers reached over the same window, plateauing at 3.9 million. Summary statistics for the time window: 19.0 million total views; 6.8 million watch hours; 105.5 million impressions (click-through rate 10.2 %; average view duration 23:04); net gain of ∼460,000 new subscribers. Data source: YouTube Studio analytics.

During the three weeks of the expedition, several posts were made on GEMPA’s Instagram profile, and its follower count increased from 900 before the expedition to more than 187,000 after. The posts captured not only the attention of the scientific community but also the general public and science students. The new followers on GEMPA’s Instagram are mainly from Argentina (92%), with smaller shares from Spain, Uruguay, Mexico, and Chile. The new followers were mostly women (73%) aged 25 to 44. In addition, during the expedition, 900 students from 18 schools across the country were connected with the scientific team throughout the Ship-to-Shore program.

During the expedition, a sea star (*Hippasteria phrygiana*), resembling the character “Patrick Star” from the cartoon SpongeBob SquarePants, was observed at ∼1000 m. The image of the sea star was chosen by the public as the expedition’s symbol and inspired numerous creative and cultural expressions. These creations included drawings, crafts, body art such as tattoos, educational materials for schools, clothing, food, etc. (**Fig. 8**). The public coined common names for several species observed during the expedition, including *estrella culona* for the seastar *Hippasteria phrygiana* (due to its buttocks-like appearance **Fig. 6D**), *batata* for the elasipodid sea cucumber *Benthodytes violeta* (“Japanese sweet potato” due to its purple coloration **Fig. 6E**), *langosta barbie* for the patagonian lobsterette *Thymops birsteini* (“barbie lobster” due to its hot pink coloration **Fig. 6F**), *merenguito* for the squat lobster *Munidopsis* sp. (“little meringue” due to its pale white coloration) and *amarillin* for an unidentified holasteroid sea urchin (due to its vivid yellow coloration).

**Figure 8.**
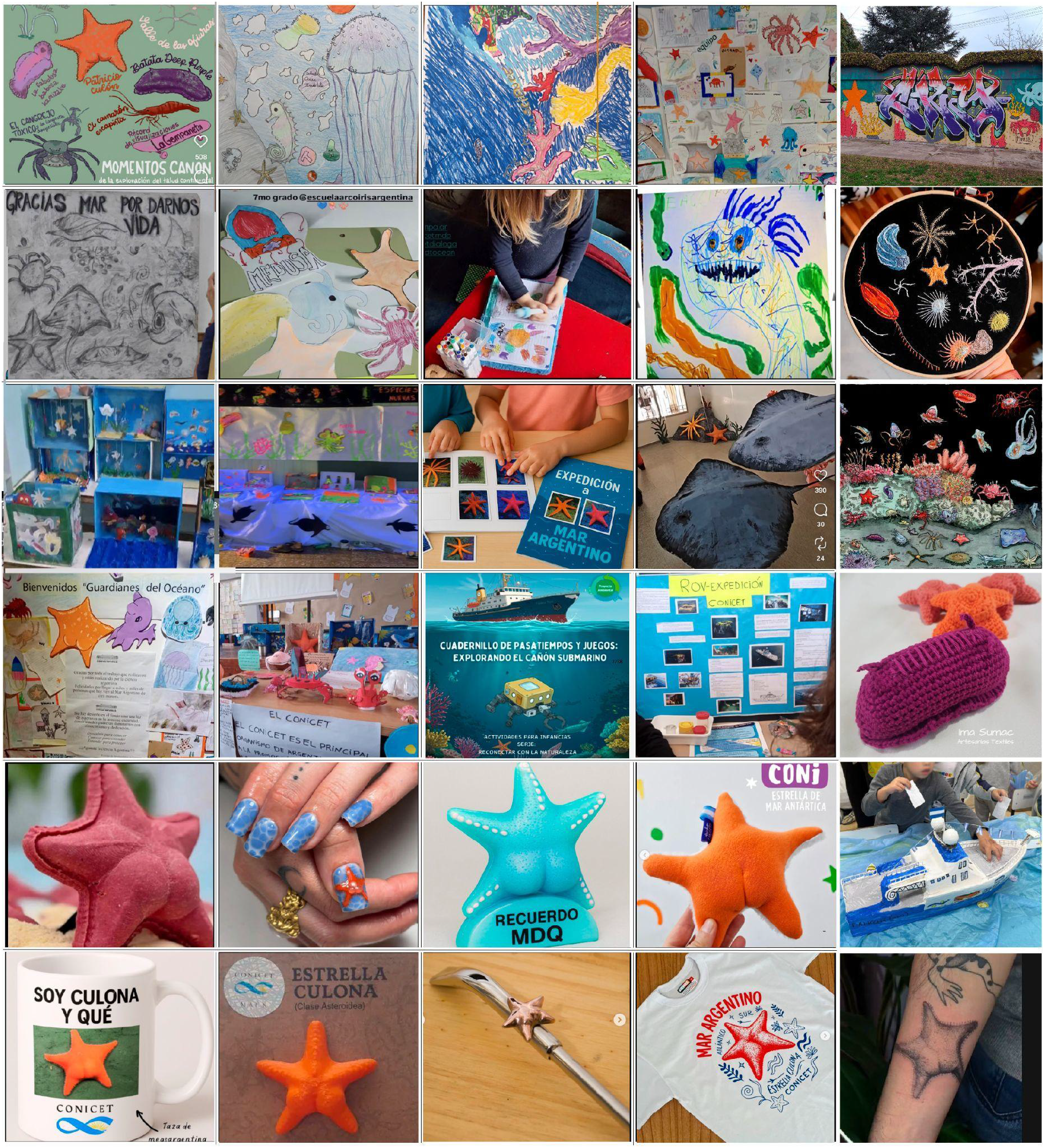
Select examples among thousands of cultural expressions that emerged during and after the expedition. Images taken from public social media posts.

During the Talud Continental IV expedition, 19 ship-to-shore sessions were organized with schools, museums, and educational institutions across Argentina and the United States, reaching approximately 900 students from 19 institutions, from kindergarten through elementary, middle, and high schools, as well as at the university level. Six participating schools (Playas Doradas, Puerto Iguazú, Barrio 31, Villa Adelina, San Andrés de Giles, and Villa Urquiza) received connectivity equipment funded by the Schmidt Ocean Institute’s small technology grants program, which remained with the institutions after the expedition. The expedition’s broad impact and visibility also led to formal recognition by multiple governmental bodies at the municipal, provincial, and national levels, including the Honorable Senate of the Argentine Nation. In total, the group and its members have received over 10 distinctions, including official commendations, declarations of legislative interest, and honorary diplomas. In addition, the expedition received awards from academic institutions (e.g., the Faculty of Exact, Physical and Natural Sciences at the National University of Córdoba), publishing and media organizations (e.g., Perfil), and the Argentine Advertising Council. Furthermore, the expedition’s live-streaming coverage was awarded the Martín Fierro Award for Best Streaming – Special Broadcast, one of Argentina’s most prestigious media awards, presented annually by the Association of Argentine Television and Radio Journalists (APTRA). Among all category winners, a single Martín Fierro de Oro (Golden Martín Fierro) is granted each year for the best overall. This distinction was awarded to the *Talud IV* expedition. The team was additionally a Webby Award nominee in the category Social & Games: Best Community or Fan Engagement. Webby Awards are the leading international awards honoring excellence on the Internet. The expedition won the Nekton Mission 2026 Ocean Literacy award.

## Conclusions

The *Talud Continental IV* expedition conducted the first ROV-based visual survey of the Mar del Plata Canyon, revealing exceptional benthic biodiversity across depths of 900–3,900 m, including extensive cold-water coral reefs, soft-coral gardens, and over 400 invertebrate species, with more than 40 species putatively new to science, six of which have already been formally described. Direct *in situ* observation via ROV provided, for the first time in Argentina, high-definition documentation of deep-sea animal distribution and behavior in natural conditions, including apparent parental defense behaviors, coordinated aggregation patterns, novel locomotion strategies, and courtship displays, underscoring the irreplaceable value of visual survey methods over trawl-based sampling for understanding deep-sea ecology and natural history.

The expedition demonstrated the scientific potential of emerging technologies in deep-sea research, including the first deployment of *in situ* eDNA filtration systems and the *DeepZoo* autonomous zooplankton sampler on ROV *SuBastian*. Anthropogenic debris, including plastics and fishing gear, was recorded at multiple depths, including sites below 3,000 m and far from the coast, confirming that even remote abyssal habitats in the Southwest Atlantic are not insulated from human influence and underscoring the urgency of integrating deep-sea ecosystems into national and international pollution and fisheries management frameworks.

The expedition achieved record-breaking public engagement for oceangoing research, demonstrating that coupling frontier deep-sea exploration with real-time, accessible outreach can generate widespread scientific awareness and inspire future generations of ocean scientists in Argentina and beyond.

Collectively, the achievements of *Talud IV* establish a critical ecological and biodiversity baseline for the Mar del Plata Canyon, providing the foundation for future deep-sea research, biogeographic analyses, and the designation of marine protected areas in the southwestern Atlantic Ocean.

## Acknowledgments

Ship-time and at-sea resources for this expedition were provided by the Schmidt Ocean Institute (expedition FKt250712). CONICET and PNA cover the Argentine scientists’ salaries. This publication is based upon work supported by CORDAP Coral Accelerator Program under Award No. CAP-2023-1502. Ocean Census, Fundación Azara, and ProyectoSub also provided funding and logistical support. JNJW received funding for expedition logistics through the National Geographic Explorer’s Program and salary through WHOI’s Access to the Sea program. DeepZoo was developed with funding from WHOI’s Innovative Technology Program and DeepTech. We are also grateful to Captain Jason Garwood and the crew of the R/V *Falkor (too)* for their strong support. Special thanks are extended to the ship’s Marine Technicians, ROV *SuBastian* technicians, and multimedia technician for their valuable assistance. Jonathan Flores helped with sea urchin determination, and Carlos Sanchez Antelo provided logistic support during the expedition.

## Notes

### Competing Interest Statement

The authors have declared no competing interest.

### Summary of Updates

Updated funding information details for CORDAP "This publication is based upon work supported by CORDAP Coral Accelerator Program under Award No. CAP-2023-1502"

